# Mechanistic corticostriatal circuit model predicts learning-dependent fMRI dynamics and individual reward bias in humans

**DOI:** 10.64898/2026.05.28.728582

**Authors:** Simon Carter, Zeming Kuang, Anthony G Chesebro, Haris Organtzidis, Sadia A Jumana, Stephen J Burke, Anand Pathak, Eva-Maria Ratai, Earl K Miller, Richard H Granger, Helmut H Strey, Lilianne R Mujica-Parodi

## Abstract

Circuit-level computational models can do more than explain existing data; they can generate novel hypotheses and capture individual differences in human populations. We demonstrate this using a biophysical corticostriatal model, transforming simulated neuronal activity from local field potential (LFP) into functional magnetic resonance imaging (fMRI) signals via the balloon model, and finally generating behavioral outcomes. We then validate the results against those obtained from human subjects. The model generates a counterintuitive yet testable prediction. Although prefrontal–striatal coherence demonstrates an *increase* at the neural LFP scale during learning, the same circuit— passed through the hemodynamic transform—–predicts a *decrease* in prefrontal–striatal BOLD correlation, driven by category-selective representations that emerge in cortex but not striatum. This prediction is confirmed in fMRI data optimized for single-subject-level detection sensitivity. The model further enables single-subject fitting of a behavioral outcome, classifying individuals into those differentially responding to positive versus negative reward bias. Established biomarkers, including activation measured by the amplitude of low-frequency fluctuations and dopaminergic effects on hemodynamic latency, are also conserved across LFP and fMRI scales. These findings reposition circuit models as generative tools for human neuroscience, capable of producing mechanistically grounded hypotheses and parsing individual variation in ways inaccessible to data-driven approaches alone.

---

Circuit-level models of neural computation have long served as interpretive frameworks for understanding animal electrophysiology, and the corticostriatal circuit mediating reinforcement learning represents one of the most thoroughly characterized examples. Dopaminergic signaling encodes reward prediction errors that drive updating of cortical and striatal representations (1), a mechanism validated across decades of non-human primate electrophysiology. Biophysically grounded circuit models built on this foundation offer mechanistic specificity and causal interpretability. They generate explicit, directional predictions, parameterize distinct behavioral phenotypes, and provide a basis for parsing individual variation in terms of underlying circuit dynamics (2). In human neuroimaging, however, such models have been largely displaced by data-driven machine learning approaches that achieve impressive performance in pattern recognition and feature extraction but are agnostic to the underlying circuitry. These approaches cannot distinguish primary neural mechanisms from secondary or epiphenomenal effects, and because they do not parameterize the generative process, they cannot produce labeled training data tied to mechanistically defined phenotypes.

While strides have been made in bridging circuit-level neural models to functional magnetic resonance imaging (fMRI) blood oxygen level-related (BOLD) signals, none have closed the loop from independently validated neural-scale dynamics to mechanistically interpretable cognitive effects in human subjects. Whole-brain spiking-network models do contain receptor-specific synaptic detail (3–5), but their interface to fMRI is benchmarked against resting-state functional connectivity, a metric that many different cellular configurations can match equally well, and they are not built around a systems-neuroscience-level circuit with neurocognitive outputs. Mean-field and neural mass reductions (6–8) reach whole-brain scale only by collapsing each region to one or two summary state variables. These models can be perturbed at an aggregate level to study circuit-level dysfunction, as in work on schizophrenia-related thalamocortical impairment (9), but they cannot resolve how a specific receptor subtype, cell class, or cortical layer would respond to an intervention. Task-specific circuit models couple biophysical neural dynamics to a forward BOLD model by summing synaptic activity, or absolute synaptic currents, and convolving with a hemodynamic response function (10–12). The neural dynamics range from spiking integrate-and-fire networks with explicit AMPA, NMDA, and GABA currents (10) to population-level firing-rate units constrained by neurophysiology (11), and the simulated BOLD reproduces qualitative features of task-evoked fMRI from the same paradigm. None of these models is independently validated against electrophysiology from the circuit and task it represents, however, so matching simulation outputs to those derived from fMRI BOLD does not by itself establish that the underlying neural dynamics are correct. Dynamic Causal Modeling (13, 14) inverts neural-to-BOLD models from data, and recent extensions replace the bilinear single-state formulation with neural-mass models that have several populations per region (15, 16), but the per-region representation is still an aggregate summary of population activity, not a circuit whose receptor- and cell-type-level dynamics have been independently validated against electrophysiology. Underlying all of these approaches, the hemodynamic interface itself remains unresolved at the cellular scale: the Balloon-Windkessel family does not specify which aspect of neural activity drives BOLD (17), and richer formulations incorporating astrocytic calcium and neuro-glio-vascular pathways (18) have thus far been shown only in single-area implementations. The result is a literature in which circuit models reproduce coarse fMRI features but have not yet been shown to deliver on what makes such models worth building in the first place: mechanistic predictions that survive translation to human data and can be used to interpret individual subjects. The stakes of closing this gap go beyond methodology. A circuit model that survives translation to human fMRI provides something that no purely statistical approach can: a generative account in which individual differences are expressed as differences in mechanistically grounded circuit parameters rather than as associations. That distinction matters for applications like predicting how a novel drug acting on a specific receptor target would alter the BOLD signal, or interpreting the cellular-scale consequences of a lesion or a developmental difference. Without such a model, fMRI in humans is restricted to description.

With one, the same imaging data becomes a powerful tool for simulating counterfactuals. With this challenge in mind, we translate a corticostriatal circuit model previously independently validated (2) against macaque electrophysiology (19) into simulated BOLD using only two transformations: the Balloon-Windkessel hemodynamic model converts the simulated neural activity into a synthetic BOLD signal, and bandpass filtering constrains it to the temporal frequencies of a real fMRI acquisition. We then compare the resulting fMRI-scale simulations against empirical 7-Tesla fMRI data from human participants engaged in category learning (Figure 1; N=31, N=62 sessions). In the Dot Category Learn Task, category prototypes are defined by spatially distinct patterns of random dots. These prototypes, unknown to participants, are inferred from ambiguous exemplars: dot patterns that are imperfect representations of each prototype. Using a forced-choice design (*Is this exemplar Category A or Category B?*), participant choices are reinforced by positive and negative feedback coupled to monetary reward (gaining money when they classify the exemplar correctly and losing money when they classify the exemplar incorrectly). Three key mechanistic signatures of the circuit survive the translation across scales. For both the simulation and human data, brain activity-based learning signatures localize to the prefrontal cortex, dopaminergic dynamics are preserved, and learning-induced PFC-striatal functional connectivity *decreases*, a circuit-based consequence of increased cortical category selectivity. This last result is particularly striking because it demonstrates how “functional connectivity,” as understood at the local field potential and fMRI scales, can fundamentally differ in sometimes counterintuitive ways. More generally, this surprising result suggests the value of a multi-scale model for generating mechanistically-derived biomarkers at each scale “bottom-up” rather than starting from fMRI-derived statistical outcomes and relying on intuitions to infer their underlying basis in terms of circuit function.

**Fig. 1.**
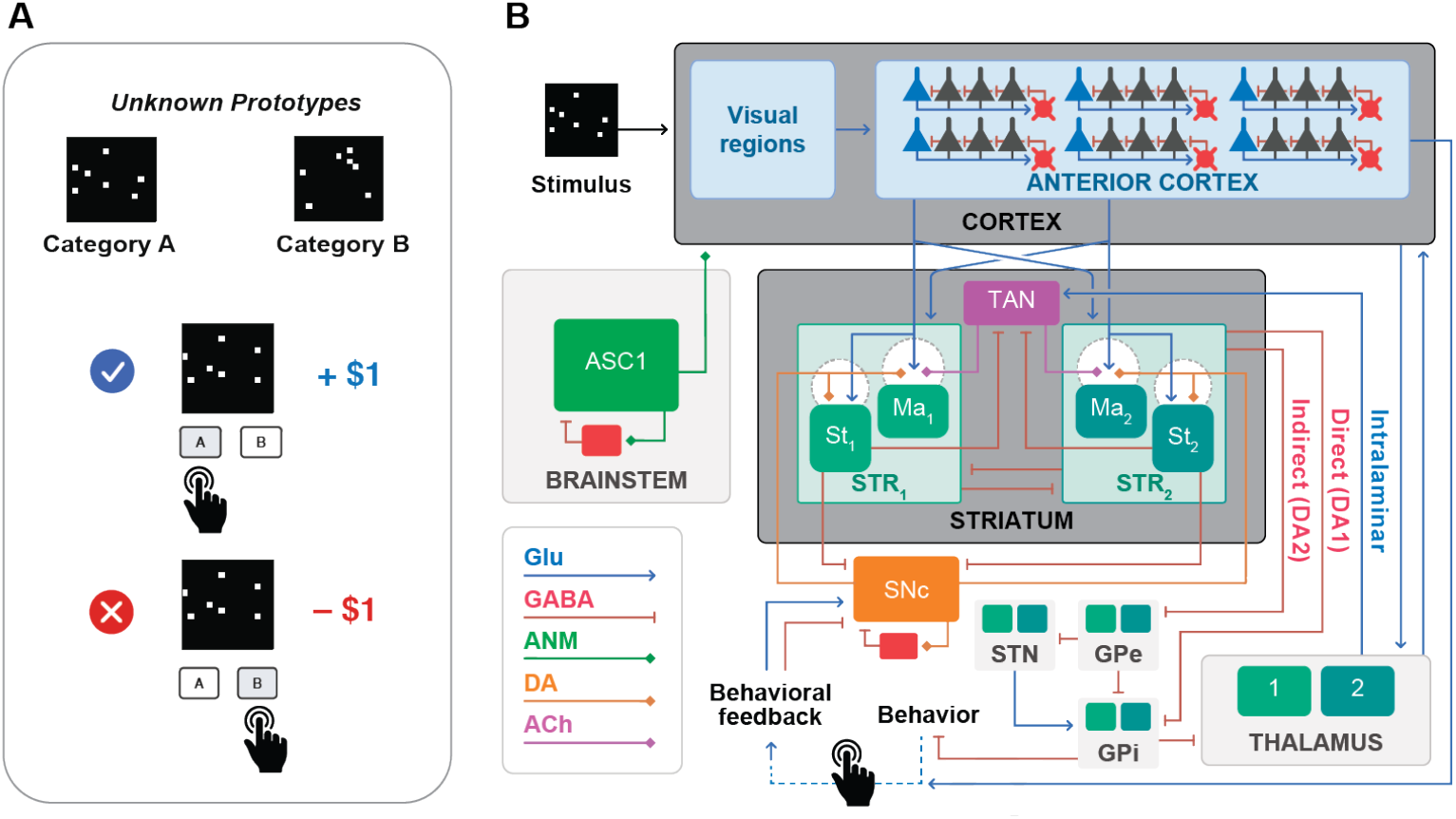
Schematic of task and circuit translated from macaque electrophysiology to human fMRI. (A) Dot Category Learn Task, in which participants infer prototype-defined categories from ambiguous exemplars, by making choices that elicit either positive or negative monetary reward. (B) Corticostriatal circuit empirically validated for the Dot Category Learn Task in macaques with reduced circuit shown shaded in grey.

Moreover, the translation enables the estimation of a behavior-predictive biomarker that would have been unattainable without a mechanistic model linking brain and behavior. Because the circuit explicitly parameterizes distinct reward-learning phenotypes, its simulation generates labeled ground-truth training data unavailable to either measured neuroimaging or behavioral data alone. Here, we demonstrate that a behavioral classifier trained entirely on simulated data stratifies human participants by reward bias, with stratification independently validated by dopamine trajectory differences that were withheld from training. To our knowledge, this is the first demonstration that a circuit model validated at the neuronal scale can be translated into human fMRI in a way that preserves its mechanistic content and yields individual-subject behavioral outputs. This proof of concept addresses a structural limitation of data-driven approaches to human neuroimaging, illustrating how biophysical circuit models can stratify human individual variation in circuit function.

## Results

### A. Conserving Dynamics: Dopamine

For the Dot Category Learn Task, the corticostriatal model’s dopamine dynamics have previously been shown (2), to start at a high concentration and decrease as learned patterns are consolidated. Specifically, the model predicts an elevated average dopamine level during early learning that converges to a baseline once participants identify the prototypes, a trajectory conserved at the fMRI scale (Figure 2A). To compare to its equivalent measured during fMRI, we used *hemodynamic speed* (HS; see Methods for details) as an empirically-established (20) surrogate for dopamine. HS demonstrates the same decrease in dopamine from the learning to the learned state (standardized regression coefficient *β* = 0.57, *p* = 0.002) (Figure 2C, D). This suggests that the dopamine-driven feedback diminishes as participants adapt to a task. While the model’s dopamine effects on synaptic plasticity do not translate to fMRI, the dopamine dynamics themselves survive. This suggests that dopamine could serve as a viable pathway for identifying and analyzing dysregulation in task adaptation. We also observed a reset of this signal when the task rule changed, consistent with dopamine’s proposed role in signaling prediction error upon environmental restructuring. Together, these results confirm that the model’s dopamine surrogate captures empirically measurable dynamics, providing a validated foundation for examining how learning-related signals are expressed across the broader corticostriatal circuit. Participants showed no significant difference in accuracy (*z* = *−*1.506, *p* = 0.1) from day 1 to day 2 (Figure S1). Hemodynamic speed (HS) demonstrated statistically significant but comparatively lower test-retest reliability (*r* = 0.45, *p* = 0.01), consistent with its sensitivity to state-dependent factors such as reward perception (Figure S2A).

**Fig. 2.**
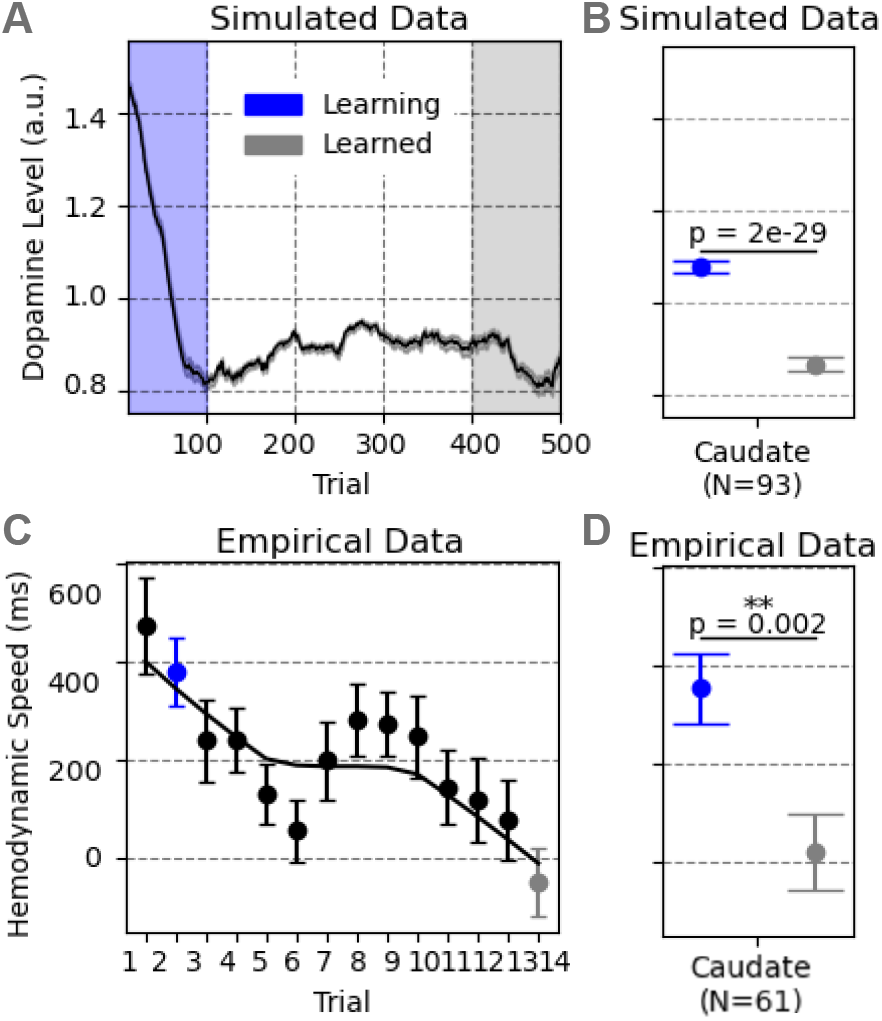
Caudate dopamine dynamics are conserved between simulated and empirical data. (A) Simulated dopamine dynamics over reward-driven learning. (B) Simulated dopamine level for the learning versus the learned state. (C) Empirical caudate dopamine dynamics over five trials, sliding window from human fMRI (N = 61 scans from N = 31 participants, including partial-cohort scans from non-completers; see Methods E for participant flow). (D) Dopamine signaling level within learning versus learned windows.

### B. Simulation-Trained Classification of Individual Reward Bias

A central challenge in computational neuroscience is translating circuit-level mechanisms into predictions about individual differences. To address this, we trained a classifier exclusively on the corticostriatal circuit model’s simulated behavioral outputs, with no access to empirical data. These model-derived signatures permitted us to partition participants into mechanistically defined reward-bias subgroups, with two parameter sets governing the magnitude of synaptic weight updates following reward (monetary gain) versus punishment (monetary loss). We then asked whether these subgroups differed on independent fMRI-derived measures that were never considered during classifier design.

Of 31 participants, 18 were classified as negative reward bias and 13 as positive reward bias (Figure 3A). Having established this partition purely from simulation, we then examined whether the groups differed in their empirical dopamine trajectories, a signal orthogonal to the classifier’s training. In the simulation, negative reward bias is associated with a monotonic dopamine decrease across learning, while positive reward bias is not (Figure 3B). Strikingly, this distinction was recovered in the empirical data without ever being sought: negative reward bias participants showed robust declines in dopamine during learning (*β* = 0.66, *p* = 0.006), while positive reward bias participants exhibited variable trajectories (*β* = 0.50, *p* = 0.07; Figure 3C). Thus, the simulation-trained behavioral classifier, applied to an independent empirical dataset, recovered subgroup differences in a neurobiological signal that played no role in its construction. This suggests that reward bias reflects a genuine, mechanistically grounded feature of dopaminergic learning dynamics, one identified by the model before it became visible in the data. Reward bias classification did not affect HS test–retest reliability in either reward class (negative class: *r* = 0.46, *p* = 0.05; positive class: *r* = 0.46, *p* = 0.1; Figure S2B).

**Fig. 3.**
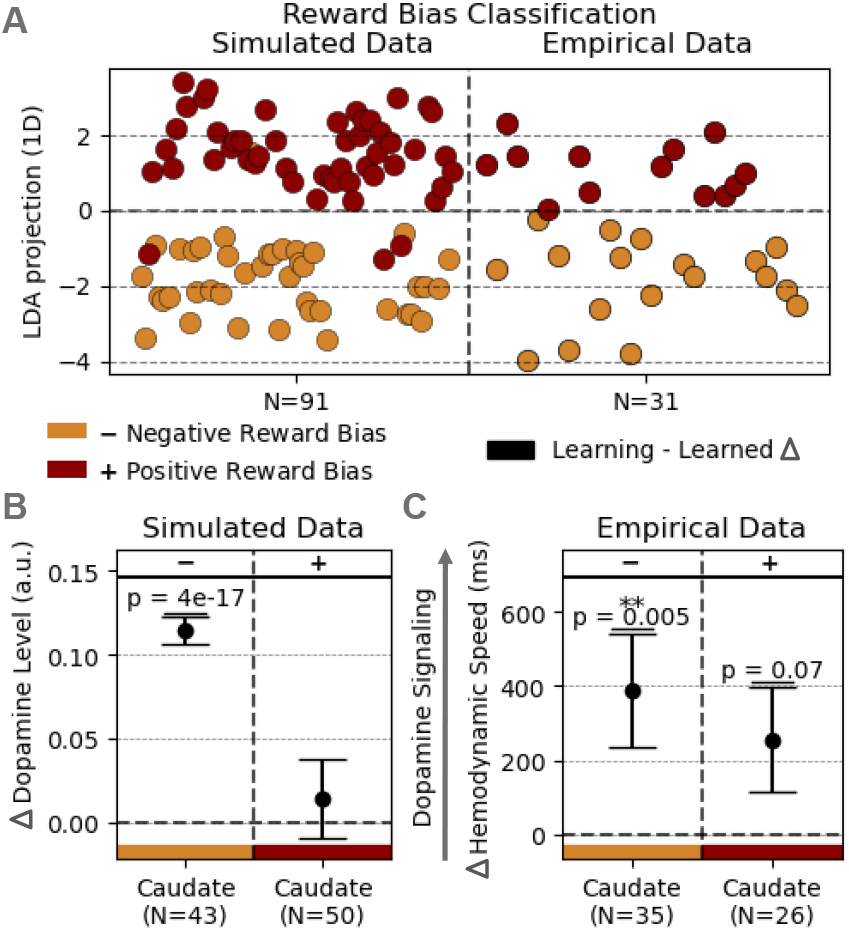
Simulation supported reward bias classification in a single subject level. (A) Behavior classification of distinct reward biases in healthy individuals. (B) Simulation indicates consistent decreases in dopamine signal in negative reward bias while inconsistent changes in dopamine signal in positive reward bias. (C) A more significant and consistent decreasing trend is shown in individuals classified as negative reward bias over the course of learning than that in positive reward bias individuals.

### C. Conserving Regions: ALFF

Beyond validating dopamine dynamics, we asked whether the model reproduces the regional pattern of neural activity changes during learning. Two questions motivated this analysis: which regions show consistency between the circuit model and empirical ALFF measurements, and given that the circuit was derived from macaque electrode recordings, how do these signals localize in the human brain? To address this, we simulated ALFF changes between the learning and learned states, finding a clear increase in anterior cortex and a smaller increase in striatum (Figure 4A). We mapped these to the human PFC and caudate, respectively, based on their functional roles and dopaminergic signatures (21–23) (Figure 4C, E, F). Empirically, ALFF increased significantly in superior frontal cortex (*β*= 0.53, *p* = 0.003) and middle frontal cortex (*β* = 0.45, *p* = 0.01), with a smaller increase in medial frontal (*β* = 0.38, *p* = 0.04), localizing the circuit’s anterior cortex signal primarily to medial and superior frontal regions (Figure 4B, D). Beyond these frontal regions, the voxelwise map also showed learning-related ALFF increases in precuneus and posterior cingulate cortex (Figure 4B); these midline regions fall outside the corticostriatal circuit modeled here and were not part of the localization target, but their engagement is consistent with their established involvement in category learning and value-based representation. Caudate ALFF showed no significant change (*β* = 0.27, *p* = 0.2), consistent with the model’s prediction of minimal striatal activity change during this transition. This convergence between simulation and imaging suggests the null caudate result reflects a genuine mechanistic property of the circuit rather than a sensitivity limitation of fMRI. ALFF showed strong test–retest reliability (*r* = 0.89, *p* = 2 *×* 10^*−*11^; *r* = 0.79, *p* = 1 *×* 10^*−*7^; *r* = 0.89, *p* = 3 *×* 10^*−*11^ in frontal regions; and *r* = 0.71, *p* = 9 *×* 10^*−*6^; *r* = 0.77, *p* = 4 *×* 10^*−*7^; *r* = 0.77, *p* = 5 *×* 10^*−*7^ in striatal regions), with the highest stability observed in regions exhibiting robust learning-related signatures (Figure S3, S4).

**Fig. 4.**
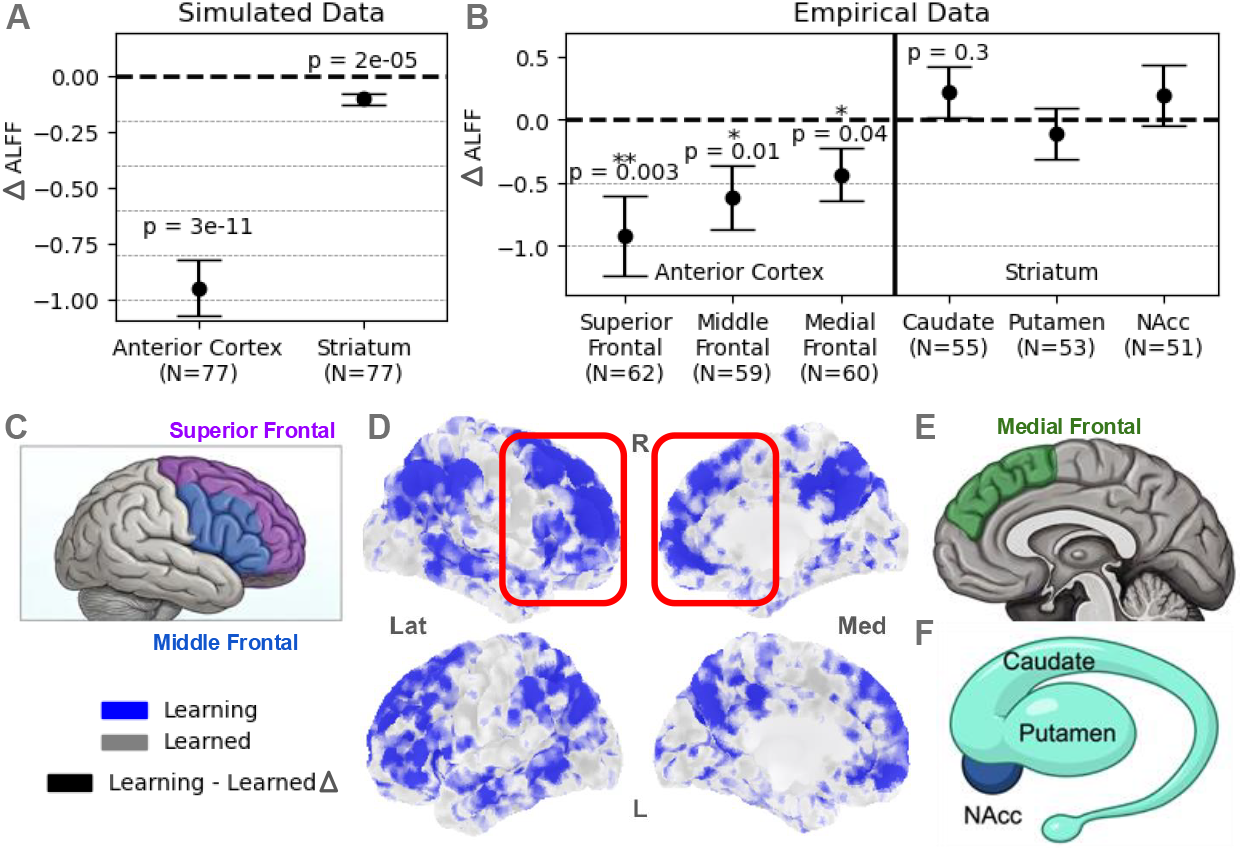
Learning-related brain activity identifies human corticostriatal equivalent of the macaque-validated circuit. (A) Simulated fMRI time-series from the human reduced corticostriatal circuit model shows differential responses in the anterior cortex and striatum. (B) Voxelwise fMRI-derived activity using ALFF localizes the learning-minus-learned signature to equivalent regions in human participants. (C) Regional quantification of ALFF learning signatures across the prefrontal cortex’s superior and middle frontal subregions, (D) shown using voxel-wise identification of extracted regions using ALFF. Other ALFF-derived regions activated by learning include the medial frontal context as shown in (E), together with (F) the striatum’s caudate, putamen, and nucleus accumbens (NAcc)

### D. Outcomes: Connectivity and Asynchrony in Brain Learning

The circuit model predicts that cortical and striatal dynamics diverge as learning proceeds. Consistent with this, ALFF increases were confined to prefrontal regions. We next tested whether this divergence extended to the functional connectivity between these structures. Comparing learning and learned states revealed a consistent, statistically significant decrease in functional connectivity between prefrontal cortex and striatum following task acquisition (Figure 5C). Across simulations, connectivity changes were heterogeneous, ranging from increases and decreases to negligible shifts. Using the empirically observed decrease in connectivity as a filter on the stochastic simulation space, we identified a subset of simulations whose dynamics matched the human data. These simulations were characterized by significantly greater and more reliable category selectivity in PFC relative to striatum, compared to simulations showing stable or increasing connectivity, motivating a formal analysis of preferential selectivity in the empirical data (Figure 5B). Following learning, category selectivity increased in PFC, particularly in medial (*β* = 0.37, *p* = 0.04) and superior frontal (*β* = 0.31, *p* = 0.08) regions, consistent with ALFF findings above and with model predictions, while striatal selectivity remained statistically unchanged (Figure 5C). Functional connectivity between these PFC regions and striatum decreased correspondingly (medial frontal–striatum: *β* = 0.43, *p* = 0.02; superior frontal–striatum: *β* = 0.35, *p* = 0.05). We propose that this reflects a shift in PFC signal composition: prior to learning, PFC and striatum share a common input driving their correlation; as learning proceeds, PFC acquires category-selective representations not present in striatum, reducing their shared signal and decreasing functional connectivity even as PFC becomes more informative. These population-level findings raise the question of whether individual variation in this corticostriatal divergence reflects meaningful differences in underlying circuit parameters, a question we address directly by examining whether simulation-derived behavioral phenotypes predict distinct neural signatures across participants (Figure 5A). As expected, preferential selectivity showed poor test–retest reliability (*r* = *−*0.14, *p* = 0.4; *r* = 0.23, *p* = 0.2; *r* =*−* 0.01, *p* = 1), reflecting variability in which stimulus category was preferentially learned across sessions (Figure S5). Functional connectivity likewise showed limited test–retest reliability (*r* = 0.14, *p* = 0.4; *r* = 0.33, *p* = 0.07; *r* = 0.30, *p* = 0.1), consistent with state-dependent learning differences (Figure S6). This dissociation from the high between-session reliability of the trait-like ALFF and hemodynamic speed metrics makes sense: a metric that tracks within-session learning-induced reorganization should not be strongly correlated across independent learning episodes from different days, whereas regional indices of stable activity properties should be. We therefore interpret functional connectivity and preferential selectivity as state markers of active learning, and ALFF and hemodynamic speed as more trait-like regional indices.

**Fig. 5.**
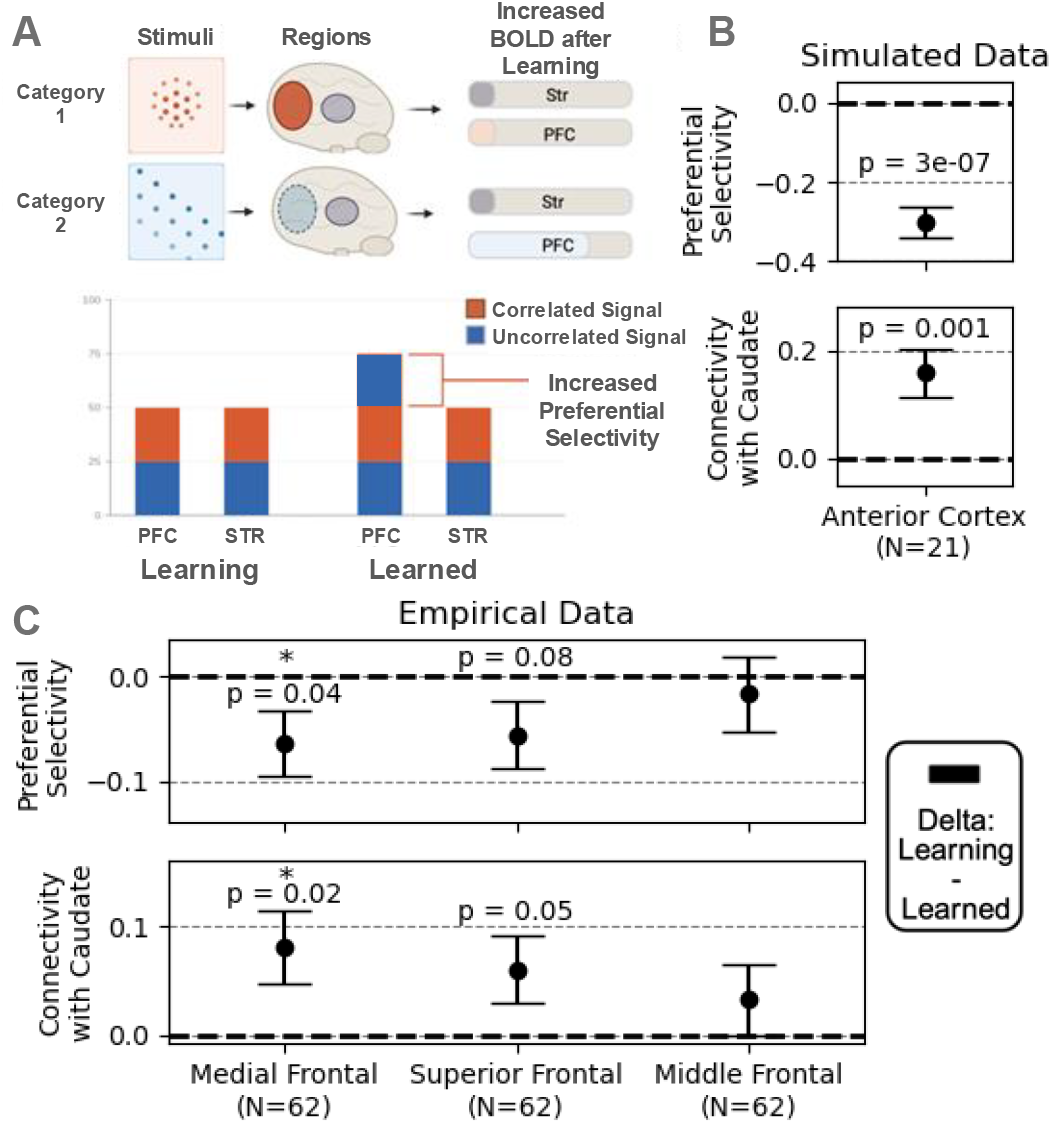
Category selectivity altered brain connectivity pattern. (A) Schematic of the preferential selectivity metric and signal mixture. (B) Simulated data showing preferential selectivity of the fMRI signal as a function of caudate–striatum correlation. Relationship between cortical asynchrony and prefrontal cortex (PFC)–striatum (STR) correlation in the corticostriatal model. (C) Corresponding metrics from empirical data.

## Discussion

Here, we present a mechanistic account of task learning grounded in a corticostriatal circuit model, validated across multiple levels of neural and behavioral measurement. Across four converging lines of evidence, we find that the model faithfully captures key signatures of human learning dynamics—and, critically, that a classifier trained exclusively on simulated behaviour successfully stratifies real individuals by reward bias, with that stratification independently validated by dopaminergic signatures never used during training.

First, we validated the model’s dopamine surrogate against empirical data. The model predicts elevated dopamine during early learning that converges to baseline once the task is acquired—a pattern confirmed in subjects, with dopamine levels showing a consistent decrease from the learning to the learned state. A reset of this signal following rule changes was also observed, consistent with dopamine’s proposed role in signaling prediction error upon environmental restructuring.

Second, examining ALFF between learning and learned states, we observed PFC activity increases consistent with circuit model predictions, localizing to medial and superior frontal cortex, with the absence of a corresponding caudate change reflecting a genuine mechanistic property of the circuit rather than a resolution limitation of fMRI. It is important to note the distinct roles played by different measures in this validation architecture. ALFF serves as the localization tool: because the macaque-derived circuit nodes carry no direct anatomical label in human space, ALFF changes are used to identify which human brain regions correspond to the simulated anterior cortex and striatal compartments. Once this localization is established, functional connectivity and hemodynamic speed provide *independent* validation of the model in those regions—they are methodologically distinct measures that were not used to define the regions of interest. The convergence of three independent signals (ALFF, connectivity, and dopamine dynamics) within the same anatomical targets therefore constitutes genuine cross-measure validation rather than circular confirmation.

Third, learning was associated with a counterintuitive decrease in PFC–striatum functional connectivity, driven by the emergence of category-selective representations in PFC that were not mirrored in striatum—a prediction generated by the model and confirmed empirically. The result is particularly striking because it runs in the opposite direction from the increase in PFC–striatum coherence reported at the LFP scale during category learning (19), a finding that the same corticostriatal model reproduces in simulation (2). A naive extrapolation from the LFP result would therefore have led the field to look for an increase in BOLD correlation in human fMRI; i.e., the *wrong feature*. Pushed through the Balloon–Windkessel transform, the model instead predicts a decrease, which is what the 7T data show, and the variance decomposition in Eqs. 1–6 makes the mechanism explicit.

That a connectivity measure can invert in sign between the neural and BOLD scales is consistent with the well-established complexity of the neural-to-hemodynamic mapping: although the BOLD signal is most closely coupled to local field potentials rather than to spiking output (24), the transformation from neural activity to BOLD is nonlinear and neurovascularly mediated rather than a simple temporal filter (25). In our model the inversion arises not from a change in the underlying coupling but from the redistribution of variance under the hemodynamic transform (Eqs. 1–6): learning-related low-frequency power added to PFC inflates its variance and lowers the BOLD correlation even as the same connectivity measure increases at the neural (LFP) scale. Consistent with reflecting the neural dynamics rather than a particular forward model, substituting a canonical hemodynamic response function for the Balloon–Windkessel transform did not substantially alter the result. Thus, our results reinforce the power of multiscale modeling while exposing the limitations of intuitively assuming a one-to-one mapping between synaptic plasticity and fMRI-derived connectivity. The spatiotemporal distinctions between scales matter not only because mechanistic detection sensitivity is lost in fMRI, but more fundamentally because they determine which fMRI signal we should look for. Moreover, one benefit of being able to pre-specify an fMRI biomarker mechanistically is that it also increases our statistical power to detect it.

Finally, and most importantly, a behavioral classifier trained entirely on simulated data successfully stratified empirical subjects into positive and negative reward bias groups. The negative reward bias group showed significantly steeper and more consistent dopamine declines—a signal orthogonal to classifier training—providing independent biological validation of the classification. Together, these results demonstrate that a mechanistically grounded corticostriatal model can account for human learning at the level of brain activity, neurochemistry, functional connectivity, and individual differences in reward processing.

### Circuit Models as Generative Tools

A central challenge in translational neuroscience is bridging the mechanistic specificity of animal models with the population-level measurements available in humans. Machine learning and AI approaches to fMRI analysis have achieved impressive performance in pattern recognition and feature extraction, but remain agnostic to the underlying biology—making it difficult to distinguish primary neural mechanisms from secondary or epiphenomenal effects, and offering little insight into the causal structure of the phenomena they characterize. By contrast, the biophysically grounded corticostriatal model used here (originally validated against macaque electrophysiology) generates explicit, testable predictions about the trajectory of dopaminergic activity, the direction and regional specificity of ALFF changes, the relationship between cortical selectivity and functional connectivity, and crucially, the behavioral signatures of distinct reward bias phenotypes. Critically, several of these predictions were counterintuitive, most notably, that learning should decrease PFC–striatum correlation due to increased cortical selectivity, and that a classifier trained purely on simulation output would recover biologically meaningful individual differences in an independent empirical dataset. Both were confirmed. The same logic informed the classifier itself. We used LDA on catch22 features so that the discriminant axis remains a linear combination of interpretable time-series statistics, and the classifier weights map directly onto the dynamical properties manipulated by the underlying circuit. This capacity to generate and validate non-obvious, mechanistically grounded predictions is precisely what distinguishes biomimetic circuit models from purely descriptive data-driven approaches. The LFP-to-fMRI transformation via the Balloon-Windkessel model serves as a principled bridge between scales, preserving key dynamical features of the circuit while enabling direct comparison with human neuroimaging. This transformation is not lossless as hemodynamic smoothing reduces temporal resolution, and regional mappings between the circuit and human anatomy involve assumptions that cannot be fully resolved, but the degree to which model predictions survive this transformation is itself an informative result. Features that replicate across the transformation are likely to reflect robust dynamical properties of the circuit rather than measurement artifacts, and speak directly to the generalizability of macaque-validated models to human populations.

### Dopamine as a Mechanistically-Grounded Biomarker

The validation of hemodynamic speed as a non-invasive surrogate for dopaminergic activity represents a practically important methodological advance. Direct measurement of dopamine in humans typically requires PET imaging with specific tracers or invasive recording approaches, both of which impose substantial constraints on experimental design and population accessibility. In contrast, hemodynamic speed can be derived from standard fMRI data, providing a scalable alternative that remains sensitive to learning-related dopaminergic dynamics predicted by circuit models.

The inference linking hemodynamic timing to dopaminergic physiology is supported by both mechanistic and empirical evidence. Dopamine modulates neurovascular coupling and vascular tone through D1- and D2-receptor–mediated effects, influencing cerebral blood flow dynamics and thereby shaping the temporal characteristics of the BOLD signal. Consistent with this mechanism, recent work has shown that voxel-wise hemodynamic latency recovers established striatal dopaminergic gradients and correlates with independent measures of dopamine function, including PET-derived indices and pharmacological manipulation (26).

Within this framework, the present caudate-specific findings provide convergent support for the validity of hemodynamic speed as a surrogate for dopaminergic signaling. The progressive decrease in signal during learning and subsequent stabilization in well-learned states are consistent with canonical dopamine dynamics associated with prediction error updating and value stabilization. The sensitivity of the metric to these structured learning phases, particularly within the caudate, aligns with its established role in integrating dopaminergic signals for adaptive learning within frontostriatal circuits.

More broadly, these findings suggest that dopamine could serve as a viable pathway for identifying dysregulation in task adaptation, even in contexts where the specific neural dynamics underlying that dysregulation are difficult to detect directly. This is particularly relevant for conditions characterized by subtle or early-stage dopaminergic disruption, where conventional biomarkers may lack sufficient sensitivity.

### Asymmetric Specialization

To formalize this mechanism, we model the regional BOLD signals as composite time series with separable components. Before learning:

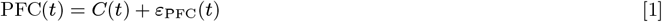

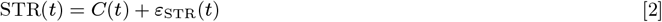

where *C*(*t*) represents common input signals (including arousal, global task engagement, dopaminergic modulation, and correlated low-frequency hemodynamic fluctuations) and *ε*(*t*) represents region-specific, uncorrelated noise. In this initial state, both regions are dominated by shared variance from *C*(*t*), resulting in high BOLD correlation (typically *r ≈* 0.85 to 0.94 in our data). After learning:

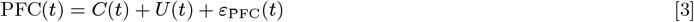

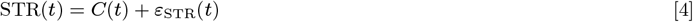

where *U* (*t*) represents learning-related activity in PFC that is uncorrelated with both *C*(*t*) and striatal activity. This component corresponds to the increase in ALFF observed in the cortex following learning: low-frequency power grows in PFC but not in the striatum, and because *U* (*t*) is uncorrelated with the striatal signal, it contributes exclusively to the uncorrelated portion of the PFC variance. After learning, while the covariance between regions remains approximately constant, the standard deviation of the PFC signal increases:

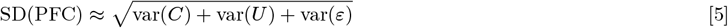

This denominator increase while the numerator stays constant results in reduced correlation (typically *r ≈* 0.45 to 0.76 in our data), proportional to the magnitude of uncorrelated low-frequency variance added to PFC. The full correlation coefficient is given by:

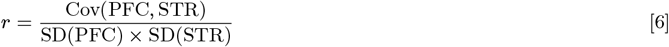

The striatum does not develop an analogous increase in uncorrelated low-frequency power, and this can be understood in terms of the specific circuit properties of the caudate. The caudate is anatomically compact, and our analyses average across two subregions whose constituent neurons, as observed in the macaque LFP data, respond preferentially to opposing categories and mutually suppress one another (23, 27–29). This opponent suppression prevents either category representation from dominating, such that averaging across subregions yields a signal that remains relatively undifferentiated throughout learning. Consistent with this interpretation, neither the model nor the empirical data showed a statistically significant increase in ALFF in the caudate, suggesting that the absence of striatal low-frequency power growth reflects a genuine mechanistic property of the circuit (specifically, the cancellation of opposing category signals) rather than a resolution limitation of fMRI. This dissociation illustrates a broader neuroscientific principle: brain regions can simultaneously undergo functional specialization and enhanced functional integration when they develop complementary roles during learning (19, 30–32). The decorrelated BOLD signals reflect the former, indicating that each region has assumed a distinct computational function. The increase in oscillatory coherence reflects the latter, enabling efficient communication between regions that now serve different but complementary roles. The prefrontal cortex develops abstract categorical representations that generalize across individual stimuli (29, 33, 34), providing top-down guidance for behavior, while the striatum maintains its role in flexible action selection through opponent encoding (27, 28). These distinct computational roles are reflected in the decorrelated hemodynamic signals observed after learning.

### Behavioral Reward Bias Reveals Distinct Dopaminergic Learning Trajectories

When participants were behaviorally classified into positive versus negative reward bias groups, distinct dopaminergic patterns emerged across learning states. In simulation, where only phasic dopamine amplitude was manipulated, the negative reward bias group showed an overall higher dopamine level compared to the positive group, along with a decreasing trend across learning. Because only phasic parameters were altered, this baseline elevation likely reflects sustained reward prediction error signaling: slower value updating maintains larger phasic bursts early in learning, which gradually diminish as contingencies stabilize (1, 35). In contrast, the positive bias group exhibited a non-monotonic (up-down) dopamine trajectory, consistent with either efficient transfer of prediction error signaling to cue-based value representations (1) or early salience amplification that normalizes with predictability (36, 37) two mechanistically distinct routes to the same behavioral outcome.

In human data, where the surrogate measure reflects caudate latency rather than amplitude (26), we do not observe a robust overall level separation between groups. Instead, the defining feature is trajectory: the negative bias group shows a clean monotonic increase in latency from early to learned states (*p* = 0.006), while the positive group shows a non-monotonic pattern that does not reach significance (*p* = 0.07). Critically, this asymmetry is precisely what the simulation predicts: the model generates a consistent, monotonic dopamine signature for negative reward bias and a variable, non-monotonic trajectory for positive reward bias. The non-significance of the positive bias result is therefore not a failure of replication but the predicted outcome—a group defined by trajectory variability should not, and does not, produce a consistent directional effect. This predicted dissociation between groups, recovered in empirical data from a classifier that never had access to neural measures, provides the key validation. We do not interpret this as evidence of the same underlying mechanism across species, but rather as convergent support for the broader claim that reward bias indexes a genuine difference in dopaminergic learning dynamics (38, 39). Notably, this distinction emerged from a classifier trained purely on simulated behavior, without ever being sought in the empirical data, suggesting it reflects a mechanistically grounded feature of striatal engagement rather than an artifact of the modeling approach.

Together, these findings point to a dissociation in how the two groups organize dopaminergic signaling across learning. Negative bias is characterized by a consistent, monotonic signature: sustained outcome-driven prediction error signaling early in learning that attenuates as contingencies stabilize (35, 40). Positive bias, by contrast, reflects a more variable trajectory whose mechanistic interpretation remains open - it may index faster cue-based value transfer (1, 40), transient salience amplification (36, 37), or heterogeneity in how individuals within this group weight recent outcomes (41, 42). This asymmetry in trajectory consistency suggests that negative reward bias may offer cleaner mechanistic traction on dopamine-related dysregulations, particularly those involving decoupling of reward prediction error from cue-based value representation (43, 44).

### Individual Differences and Clinical Translation

Beyond group-level validation, the model supports single-subject classification, distinguishing individuals according to positive versus negative reward bias based on behavioral data alone. The subsequent observation that these behaviorally derived groups differ systematically in their dopamine trajectories, a signal not used during classifier training, provides independent validation of the classification and suggests that reward bias captures a meaningful axis of individual variation in the underlying circuit dynamics. The convergence of this pattern across both simulated and empirical data further strengthens confidence in the classifier’s validity. The ability to distinguish between positive and negative reward bias carries direct clinical resonance for mood disorders: relative insensitivity to reward, coupled with heightened sensitivity to punishment, characterizes anhedonia in major depression (45, 46), whereas exaggerated reward sensitivity is a hallmark of the (hypo)manic pole of bipolar disorder (47). A behaviorally derived classifier grounded in a mechanistic model could offer a non-invasive means of stratifying participants (48), tracking treatment response (49), or identifying individuals at risk before symptoms become clinically apparent (50). Although the present study is limited to a healthy population engaged in a controlled learning task, the framework is in principle extensible to clinical cohorts.

### Limitations

Several limitations of the scaling procedure warrant explicit acknowledgment. First, the LFP-to-fMRI transformation via the Balloon model (51) introduces hemodynamic smoothing that reduces temporal resolution, potentially obscuring fast dynamics present in the original circuit. Second, the mapping between circuit regions and human anatomical structures involves assumptions that cannot be fully resolved; in particular, the cortical region of the circuit encompasses both anterior and visual cortex, and its localization in the human brain remains ambiguous. Third, the model was originally derived from electrode recordings in macaques, and between-species differences in cortical geometry and connectivity (52) may limit the precision of regional predictions in humans. Fourth, the lower task resolution and comparatively slower learning dynamics in the simulation necessitate caution when drawing direct comparisons to human behavioral and neural data (53), as these discrepancies may introduce systematic differences that are not easily dissociated from genuine between-species variation (54). Fifth, the current implementation does not model transitions between distinct task contexts (55), constraining the generalizability of the findings to settings in which cognitive demands remain relatively stable. Finally, the present sample of 31 subjects, while sufficient for the analyses reported here, limits the generalizability of the single-subject classification results (56). Future work with larger and more diverse samples will be necessary to establish the robustness of the reward bias classifier across populations (57). These limitations should be understood not as deficiencies of the approach but as principled boundaries on interpretation. The degree to which model predictions survive these constraints and replicate in human data is itself a strong positive result, suggesting that the core dynamical properties of the corticostriatal circuit are preserved across species and measurement modalities (58).

### Future Directions

Several natural extensions of this work suggest themselves. Most immediately, applying the classification framework to clinical populations—including individuals with Parkinson’s disease, schizophrenia, or substance use disorders— would test whether reward bias and its associated dopaminergic signature are altered in conditions known to involve corticostriatal dysfunction. Longitudinal application of the hemodynamic speed measure could enable tracking of dopaminergic changes over the course of learning or treatment, providing a dynamic biomarker sensitive to individual trajectories rather than cross-sectional snapshots. The scaling procedure developed here could in principle also be applied to other validated circuit models, extending the generative approach to different cognitive domains and neural systems. Finally, integrating the behavioral classifier with additional imaging modalities—such as PET or MRS—could provide convergent validation of the dopaminergic signal and refine the mechanistic interpretation of individual differences in reward bias.

## Methods

### E. Participants and clinical assessment

Functional MRI data were acquired during a reward-driven learning task from a cohort of healthy adults (N = 31; mean age = 30.33 *±*7.29 years; 12 females) aged 18–45 years to investigate learning-related neural dynamics in the human brain. The study was registered on ClinicalTrials.gov (identifier: NCT06373016) and approved by the Institutional Review Board (IRB) at Massachusetts General Hospital (Boston, MA), with additional approval from the IRB at the State University of New York at Stony Brook (Stony Brook, NY).

Participants were recruited from the Boston metropolitan area through advertisements. Exclusion criteria included MRI contraindications, neurological or psychiatric disorders, history of brain injury, recent recreational drug use, or heavy alcohol consumption.

All participants provided written informed consent prior to participation. Participants underwent a physical examination and completed the Standardized Mini-Mental State Examination (SMMSE) to confirm cognitive normalcy. All 31 enrolled participants completed both scanning sessions and formed the primary analysis cohort, yielding 56 sessions. Dopamine-trajectory analyses presented in Figure 2C additionally include partial-cohort scans contributed by the enrolled subjects who did not complete the longitudinal protocol, for a total of N = 62 scans from N = 31 unique participants. Although motion-corrupted scans were excluded (Methods H), region-specific analyses (ALFF, functional connectivity, and category selectivity; Figures 4 and 5) could draw on the entire cohort for individual unaffected components. These yielded region-specific sample sizes of up to N = 62, which also varied by region due to IQR outlier removal (Methods N).

### F. fMRI Acquisition

Brain images were acquired at an ultra-high-field (7T) MRI scanner at the Massachusetts General Hospital Athinoula A. Martinos Center for Biomedical Imaging. Each participant follows the same protocol twice on two separate days. The sequences include a brain localization, a 1 mm isotropic T1-weighted structural (multi-echo magnetization prepared rapid gradient echo [MEMPRAGE]) image, and whole-brain BOLD (echoplanar imaging, EPI) for the task session. BOLD images were acquired using the following protocol: Simultaneous multi-slice (SMS) slice acceleration factor = 5, R = 2 acceleration in the primary phase encoding direction (62 reference lines) and online generalized autocalibrating partially parallel acquisition (GRAPPA) image reconstruction, TR = 802 ms, echo time (TE) = 20 ms, flip angle = 33°, voxel size = 1.75 × 1.75 × 1.75 mm, slices = 85, and number of measurements = 780. The whole-brain T1-weighted structural volumes were acquired with 1 mm isotropic voxel size and four echoes with the following protocol parameters: TE1 = 1.61 ms, TE2 = 3.47 ms, TE3 = 5.33 ms, TE4 = 7.19 ms, TR = 2,530 ms, and flip angle = 7°, with R = 2 acceleration in the primary phase encoding direction (32 reference lines).

### G. Dot Category Learn Task

The task was adapted from a paradigm extensively used in cognitive neuroscience and previously implemented in macaques (19). Participants were instructed to categorize dot-pattern images into one of two categories without prior knowledge of the underlying classification rule. Within each stimulus set, two prototype images corresponding to the two categories were pre-defined but not shown. Instead, participants were presented with distorted versions of these prototypes, from which they had to infer categories from imperfect exemplars. Across 20 trials per stimulus set, participants were expected to gradually infer categories based on their choices and trial-by-trial monetary feedback, which consisted of either positive or negative reward.

Each fMRI run consisted of four consecutive stimulus sets. On each trial, a stimulus was presented for up to 2.3 s, during which participants were required to indicate the category assignment before stimulus offset. Correct responses were rewarded with$1, whereas incorrect responses or failure to respond within the allotted time resulted in a$1 loss. Feedback indicating the monetary outcome was presented immediately after each decision. An inter-trial message then appeared for up to 1.6 s, which participants could terminate early to proceed to the next trial.

### H. fMRI Preprocessing

For MRI images, subject-level T1w and fMRI images were preprocessed using fMRIPrep (https://fmriprep.org/), version 20.2.3 (59). Following fMRIPrep, denoising was performed using BrainDancer (ALA Scientific) cleaning implemented via BDtools (60) (https://github.com/hstrey/BDTools.jl). Confound removal, bandpass filtering (0.01 Hz to 0.1 Hz for ALFF and FC; 0.009 Hz to 0.15 Hz for hemodynamic speed (26)), detrending, and standardization were performed using Nilearn (https://nilearn.github.io/). Framewise displacement (FD) was computed for each scan. Scans with a mean framewise displacement (FD) greater than 0.5 mm were excluded from analysis. All scans included in this study met this criterion. White matter, CSF, and motion parameters (three translations and three rotations) were included as confound regressors. Parcellation from whole brain voxel space to region space follows Talairach definition of gray matter regions (61).

### I. Scaling-Up Procedure

A schematic of the full scaling-up pipeline is shown in Figure 6. We begin with the previously validated corticostriatal circuit model (2), which outputs at the level of spiking neurons, synaptic plasticity, local field potentials (LFPs) and behavior, predicting electrophysiological and learning responses in the rhesus macaque (19). To bridge from this mesoscale electrophysiological signal to the Blood-Oxygen-Level-Dependent (BOLD) signals measured by fMRI, we apply the Balloon-Windkessel model (51, 62) to the simulated LFP output. Balloon model parameters follow the experimentally derived values from (62), which attenuate high-frequency content more strongly than the theoretically derived parameter set from the same paper. This behavior produces a closer match to the standard HRF and emphasizes the frequency range considered neuronal in the fMRI literature (63). This yields a simulated hemodynamic response function for each region in the circuit. To facilitate comparison with empirical fMRI data, we apply a 5th-order Butterworth bandpass filter retaining frequencies in the 0.01–0.1 Hz range, consistent with the conventional resting-state fMRI band (63). We verified that replacing the Balloon-Windkessel model with a standard canonical HRF convolution yielded no substantial differences in the resulting timeseries, supporting the robustness of our approach to this modeling choice.

**Fig. 6.**
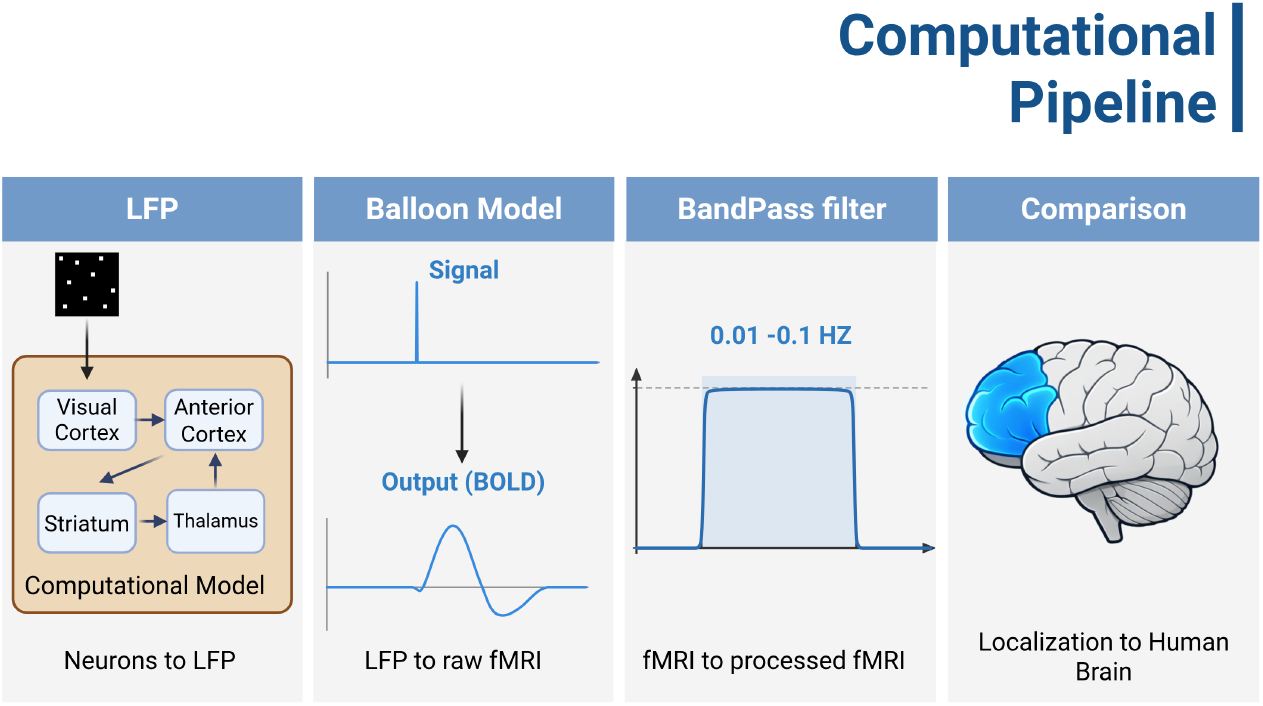
Schematic of the scaling-up pipeline. The simulated LFP output from the corticostriatal circuit model is passed through the Balloon-Windkessel model to generate a hemodynamic signal, which is then bandpass filtered (0.01–0.1 Hz, 5th-order Butterworth) to produce simulated fMRI timeseries. Finally, ALFF-based biomarkers are used to localize the abstract circuit nodes to anatomical regions of the human brain.

### J. Hemodynamic Speed, a Time-Varying surrogate for Dopamine Dynamics

Dopamine dynamics from the human fMRI were approximated using a previously validated hemodynamic latency metric (26). For each voxel, relative latency was computed with respect to the global brain signal by identifying the temporal shift that maximized the cross-correlation between the voxel time series and the global signal. To capture task-dependent state changes (e.g., learning vs. learned phases), fMRI time series were segmented into sliding windows comprising five consecutive trials. Latency estimates were computed within each window to characterize dynamic changes across task states. In the original method, hemodynamic latency was inversely associated with putative dopamine signals. To aid in interpretability, we reversed the metric’s sign such that higher values (hemodynamic 4 speed) correspond to stronger dopamine-related effects.

### K. Amplitude of low frequency fluctuation (ALFF)

ALFF reproduced learning signatures in empirical data and localized the simulated anterior cortex in hypothesized human brain regions. Due to the nature of the corticostriatal circuit and its validation in macaques, localization of individual simulated brain compartments was not obvious and required validation. ALFF and functional connectivity are the most published metrics in fMRI. While functional connectivity is an edge level metric to infer connectivity between regions, ALFF provides an indication of changes in neuronal signal variation for each brain region. Defined as the variance of the signal after bandpass filtered into the 0.01 to 0.1 Hz range, the ALFF was computed at the voxel level for each fMRI scan. The time axis was windowed the same way as the dopamine measures for both empirical and simulated data, splitting early (learning; mean accuracy: 0.630) from late (learned; mean accuracy: 0.746) task states. For empirical data, voxel-level ALFF values were normalized to the average white matter ALFF within each time window to remove background noise. While simulated data produced two ALFF values per simulation (early versus late; *z* = 4.935, *p* = 8*e*^*−*7^), a within-scan paired t-test was performed at the voxel level to identify significant voxels in the empirical data. The empirical data were then parcellated into regions of interest following the gray matter regions defined in the Talairach atlas. Regions highlighted in the localization step were subsequently compared using a within-scan paired t-test.

### L. Category Selectivity and Functional Connectivity Analysis

Forty three stochastic simulations of the corticostriatal model were computed. In each run, the binary pulse train was concatenated five times to match the empirical timeseries length, accounting for the 8 second per trial structure of the task relative to the 1.5 second simulator resolution prior to application of the BOLD model. Category selectivity was quantified by computing the cross-correlation between a raw binary pulse train representing the stimulus presentation (1 for category A, 0 for category B) and the BOLD signal in each region of interest. Each pulse was made the same length as each trial. The cross-correlation was computed over a 0–10 second lag window, and the magnitude of the first peak was used as the selectivity metric. The window was restricted to 10 seconds to avoid ambiguity from oscillatory peaks at longer lags, while empirically this was done for 16 seconds. Higher values reflect stronger differential responses between categories independent of response direction or hemodynamic delay. Functional connectivity between PFC subregions and striatum was computed as the Pearson correlation between regional mean BOLD timeseries within learning and learned state windows respectively, as defined above. Group-level differences in connectivity and selectivity between states were assessed as described in the Statistics Section. To identify simulation regimes consistent with the empirical data, simulations were filtered by the direction of their PFC–striatum connectivity change, retaining those showing a decrease from the learning to the learned state. Of the 43 simulations, 26 exhibited category selectivity during the learning state (i.e., the peak detection algorithm produced valid results). In addition to exhibiting category selectivity, 25 of these 26 simulations also showed higher category selectivity in the learning compared to the learned state. Category selectivity was then compared between the filtered subset and the remaining simulations, as described in the Statistics section. The filtered subset was taken as the biologically plausible regime consistent with human data.

### M. Classifier

We trained a classifier to distinguish positive and negative reward bias phenotypes in both the Model and patient scans, undergoing the same task. Model parameters were selected in accordance with (2). Each phenotype was simulated across multiple runs. As classifier accuracy showed minimal improvement beyond the first 500 trials, with no appreciable gains observed in the final 200, only the initial 500 trials per run were retained for analysis. To account for behavioral differences and align with empirical scan resolution (given that humans acquire dot pattern associations considerably faster) the 500 trials were windowed into bins of 20. The windowed data were then projected into catch22 space (64), from which a Linear Discriminant Analysis (LDA) classifier was trained to differentiate the two phenotypes: 41 positive reward bias and 50 negative reward bias. We used LDA rather than a nonlinear classifier (e.g., random forest, gradient-boosted trees, or a neural network) because the LDA discriminant axis is a linear combination of the input catch22 features. Each catch22 feature is itself a defined time-series statistic (64)—autocorrelation structure, distributional shape, stationarity, entropy, and so on—so a linear projection preserves the mapping from classifier weights back to these statistics. A nonlinear classifier might achieve higher accuracy, but would obscure which features of the trial-by-trial behavior separate the two phenotypes. Since the broader rationale for using a biophysically grounded circuit model is mechanistic interpretability, we preferred a classifier whose decisions can be read back in terms of identifiable behavioral statistics. For analysis and validation, we then looked at the dopamine levels learning vs. learned in the participants.

### N. Statistics

All statistical analyses were conducted using linear mixed-effects models to account for the repeated-measures structure of the data. Each participant completed two scanning sessions, and both sessions were included in the analysis. For each metric, outliers were identified and removed using the interquartile range (IQR) method, with observations exceeding 1.5 × IQR beyond the first or third quartile excluded prior to model fitting. Group-level differences between early and late task phases were assessed using mixed-effects models of the form metric *∼* State (early vs. late), with State specified as a fixed effect and subject included as a random intercept to account for within-subject dependency across sessions. Effect sizes for fixed effects were quantified using standardized regression coefficients (*β*), calculated by dividing the estimated fixed-effect coefficient by the residual standard deviation of the model, providing a Cohen’s d–equivalent measure for interpretability. Statistical significance was evaluated using two-tailed tests with an alpha level of 0.05. Test–retest reliability was assessed at the session level by first computing the mean value of each metric within each session for each subject. Reliability was then quantified as the Pearson correlation coefficient between session 1 and session 2 across subjects, providing a measure of between-subject consistency across sessions.

## Data Availability

The model presented in this work is part of the Neuroblox computational neuroscience platform (code: https://neuroblox.ai/code; documentation: https://neuroblox.ai/docs). Following publication, all data, tutorials, and a standalone implementation to reproduce the results of this work will be available at https://neuroblox.ai/pubs.

## Author Contributions

L.R.M.P. conceived and supervised the study, acquired funding, and wrote the manuscript. S.C. developed the computational modeling, scaling-up procedure, and techniques for reward-bias classification and category selectivity. Z.K. analyzed the fMRI data, conceived of the dopaminergic analyses, and wrote the manuscript. H.O. developed the task translation from macaque to human, and developed and analyzed the behavioral modeling. A.P., R.H.G., and E.K.M. assisted with the corticostriatal scaling-up procedure. H.H.S. assisted with the parameter estimation and software implementation. E.M.R., A.G.C., S.A.J., and S.J.B. contributed to data acquisition. All authors reviewed, edited, and approved the final manuscript.

## ACKNOWLEDGMENTS

The research presented here was funded by the Baszucki Foundation (to LRMP), using NIH-supported resources from the Massachusetts General Hospital Translational Clinical Research Center (1UL1TR002541-01). AGC also acknowledges support from NIHGM MSTP Training Award (T32-GM008444).

