## Supplemental Figures for "Mechanistic corticostriatal circuit model predicts learning-dependent fMRI dynamics and individual reward bias in humans" (from another version of this preprint: version 2, 2026-06-08)

S1

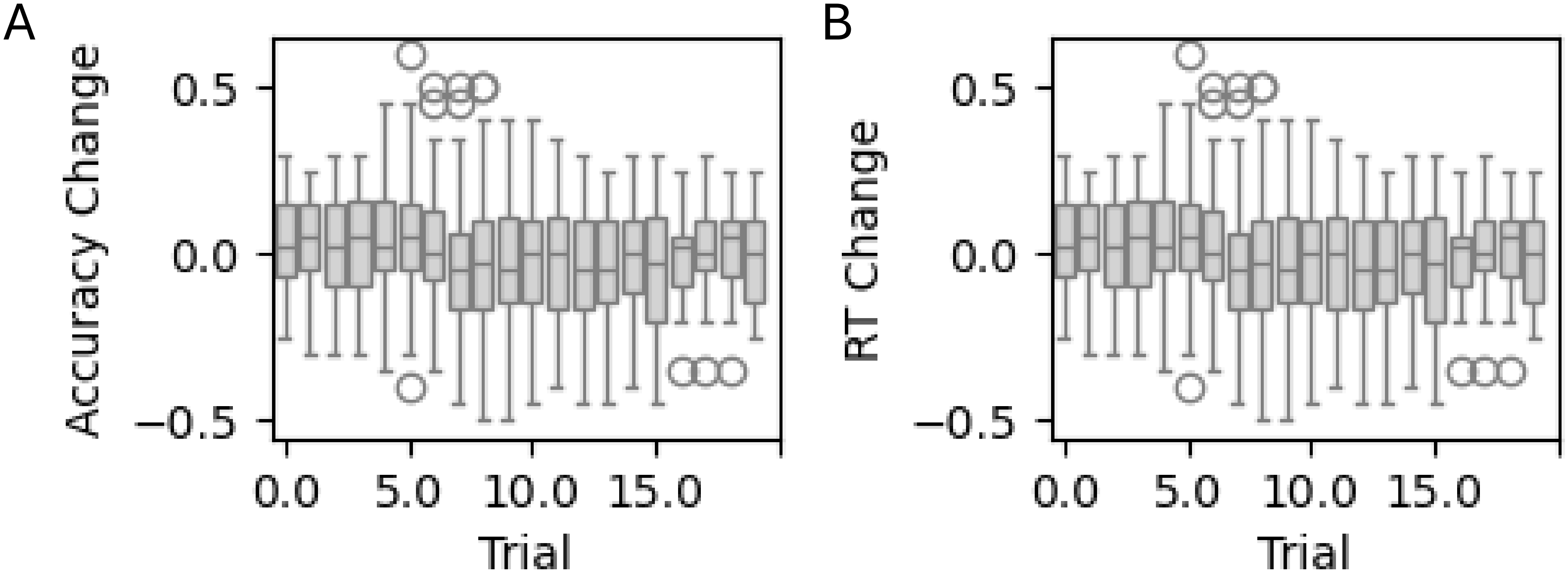


Within subject performance consistency scan sessions. A) Delta accuracy and B) delta response time

S2

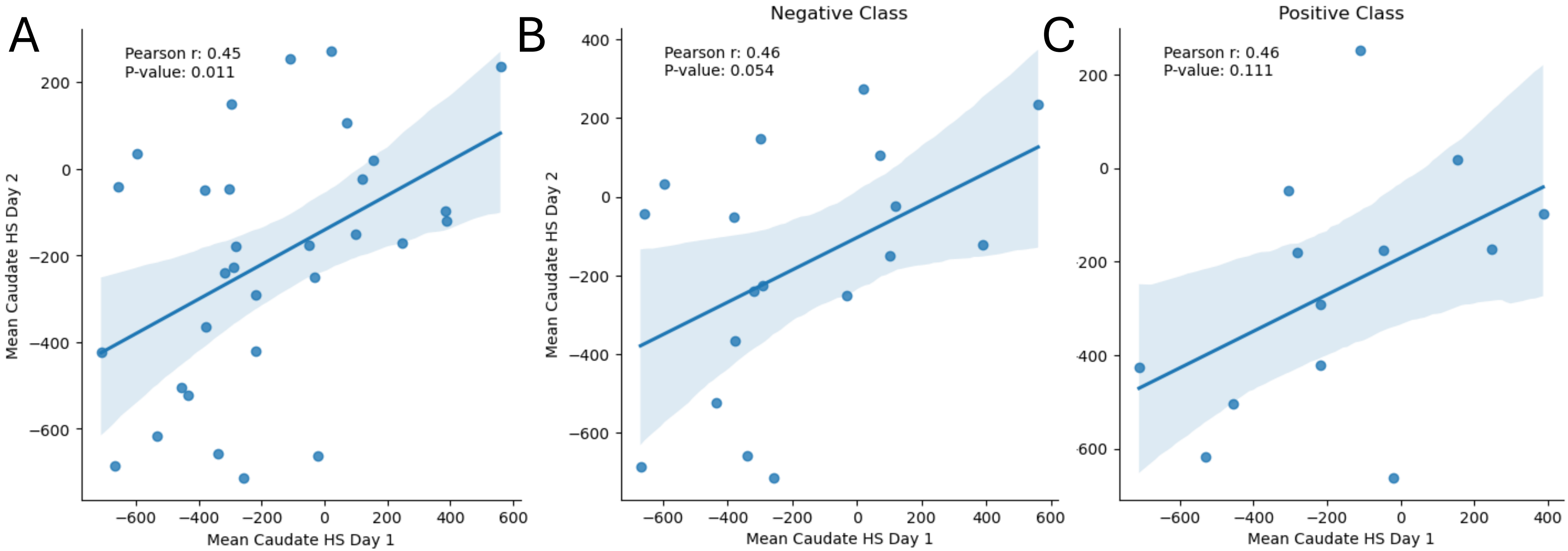


Test-retest reliability of hemodynamic speed across scan sessions. A) Whole cohort reliability, B) negative reward bias class reliability, and C) positive reward bias class reliability

## S3

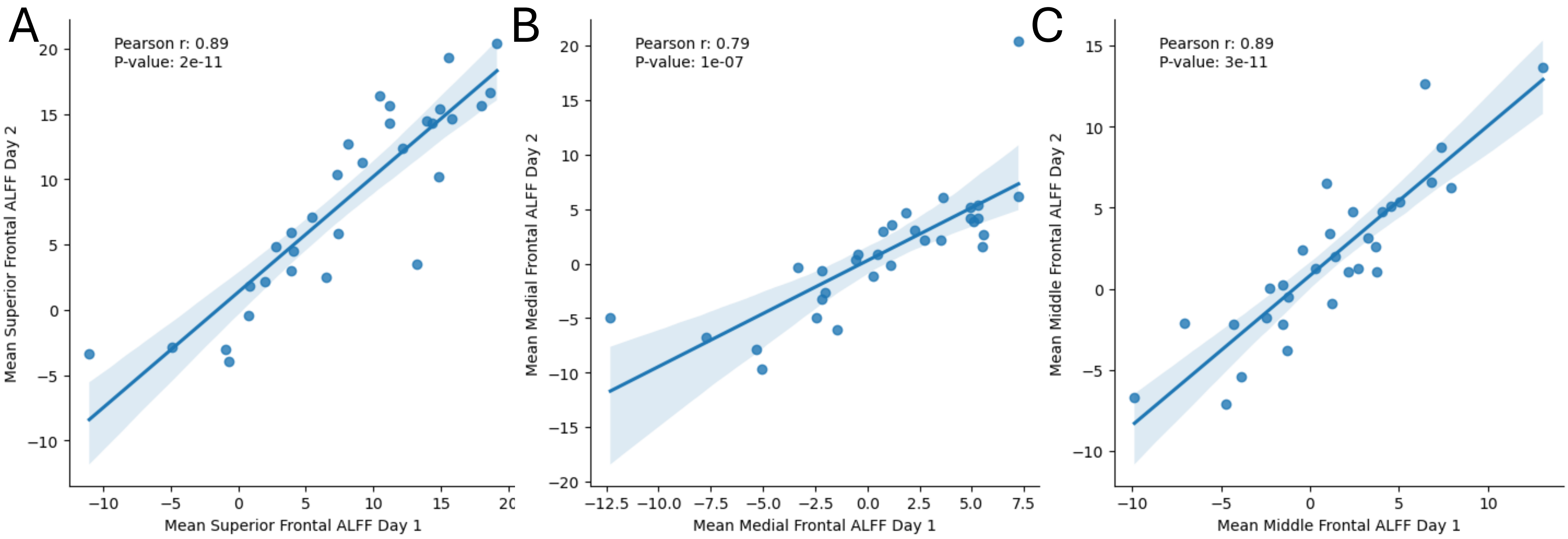


Test-retest reliability of ALFF in frontal areas across scan sessions. A) Superior frontal, B) medial frontal, and C) middle frontal.

# S4

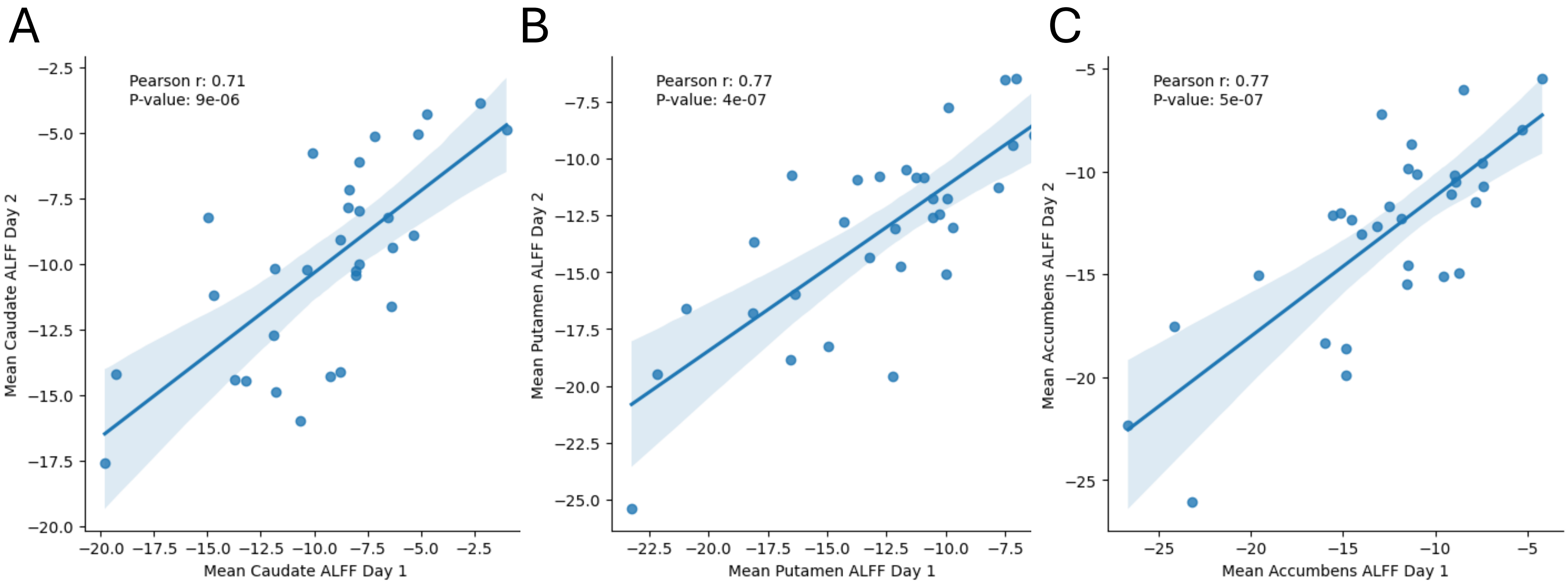


Test-retest reliability of ALFF in frontal areas across scan sessions. A) Caudate, B) accumbens, and C) putamen.

S5

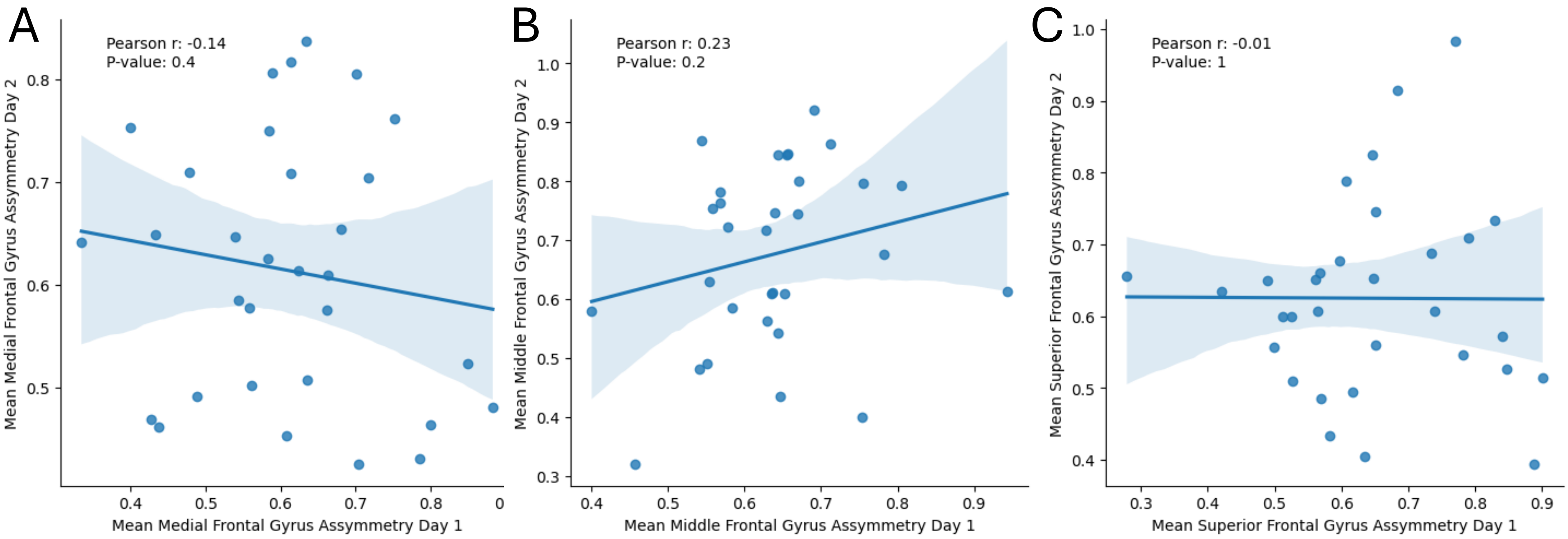


Test-retest reliability of preferential selectivity (asymmetry) in frontal areas across scan sessions. A) Medial frontal, B) middle frontal, and C) superior frontal.

S6

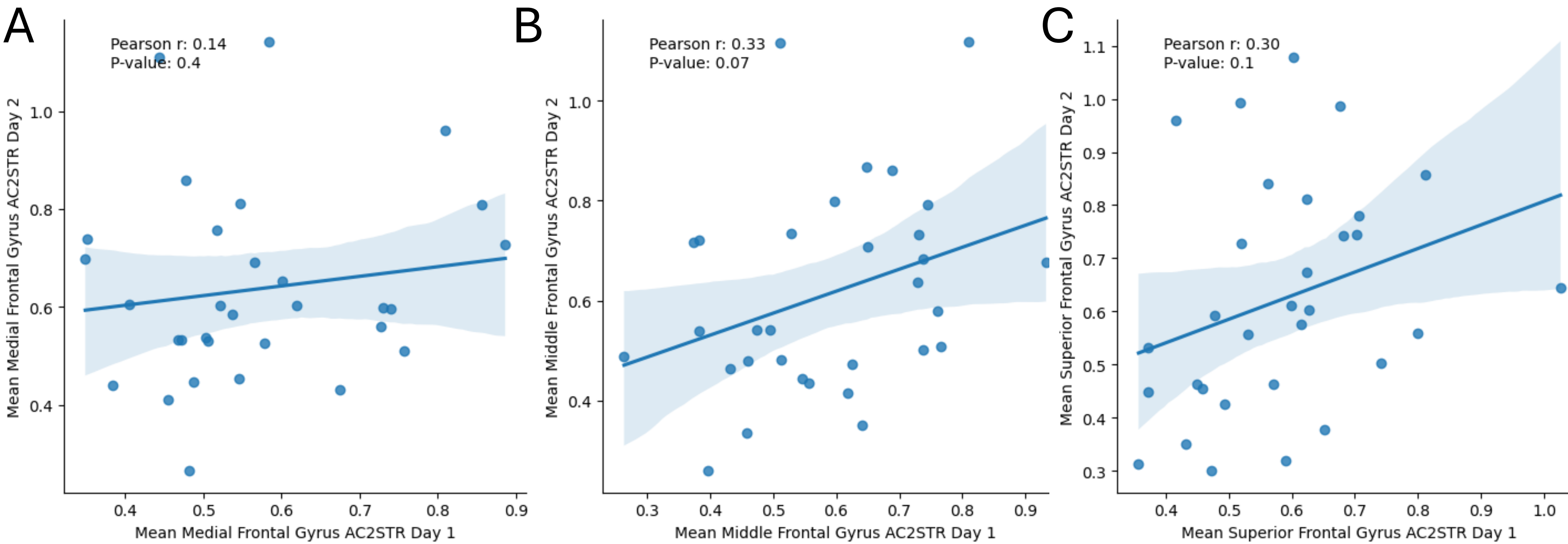


Test-retest reliability of functional connectivity with Caudate (AC2STR) in frontal areas across scan sessions. A) Medial frontal, B) middle frontal, and C) superior frontal.
